# Natural Selection in the Wake of Catastrophe

**DOI:** 10.64898/2026.06.08.730870

**Authors:** Jesse Young Lin, Omer Granek, Joshua Sodicoff, Seppe Kuehn, David Pincus, Vincenzo Vitelli

## Abstract

Living organisms, from bacteria to humans, are more likely to survive if their traits enhance fitness. In populations well adapted to their environmental niches, natural selection proceeds via rarely beneficial mutations. But when a catastrophe wipes out niche diversity, sudden adaptation often follows. Here, we present a data-validated theory of natural selection in the wake of catastrophe and unveil a simple law that emerges during recovery: the mean fitness relaxes inversely with time, with a prefactor proportional to the number of traits coupled to the post-catastrophe environment. We put our approach to test using experimental fitness landscapes measured following antibiotic administration to *E. coli*. The resulting mean trait adaptation is not described by gradient ascent on a fitness landscape, instead it follows an algorithm known as Levenberg–Marquardt optimization. Near fitness peaks, evolutionary trajectories are biased against greediness — from an optimization perspective, post-catastrophic selection is optimistic.

---

Recovery from catastrophe is a collective effort that shapes the evolution of complex systems from human societies to ecosystems. After an ecosystem collapse (Fig. 1(a)), a mutagenic stress (Fig. 1(b)), an acute clinical event [1, 2], or a natural hazard [3], the cohort of affected individuals, be it bacteria or humans, begins its recovery at time *t* = 0. What follows is rarely governed by a single, typical timescale: the aggregate response reflects a heterogeneous mix of units with latent “traits” (i.e. covariates) **z** = (*z*_1_, …, *z*_*n*_), each failing or rebounding at its own characteristic rate *r*(**z**) [4–8]. For instance, following a layoff, an individual’s characteristic time to re-employment 1*/r*(**z**) depends on traits such as individual-specific wage offers or unemployment-benefit duration [9–11], while other unmeasured traits are subsumed into a random “frailty” [10, 12]. The recovery itself is selective: traits with high *r*(**z**) come to dominate the aggregate response [4, 13]. Yet in many settings, the origin of this trait heterogeneity is difficult to disentangle from the complexity of the underlying dynamics [6, 10, 12, 14]. Here, we ask: what is the average recovery trajectory in the aftermath of catastrophe?

**FIG. 1.**
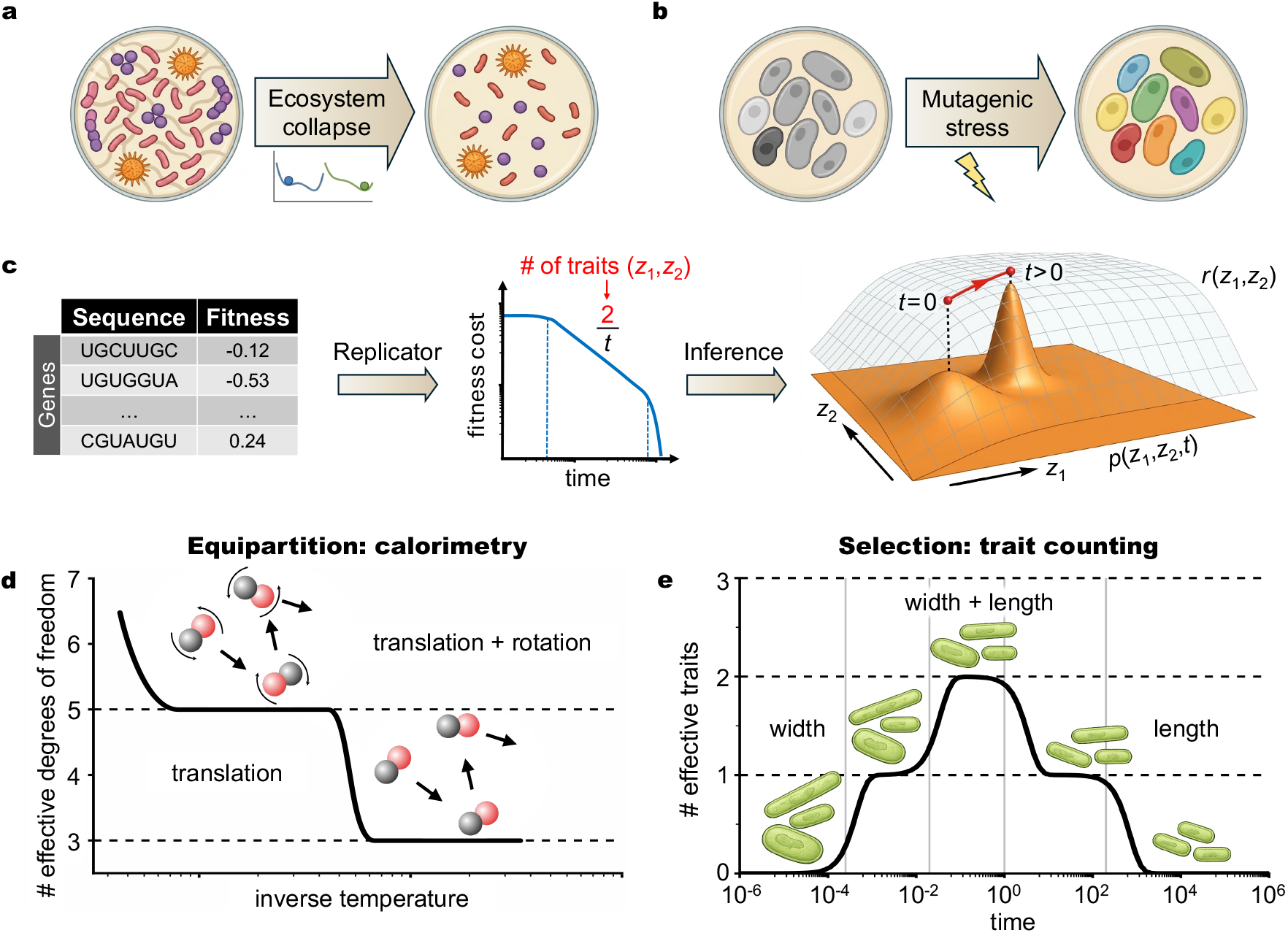
Catastrophes turn standing variation into a dynamical probe of fitness-landscape geometry. **(a)** Ecosystem collapse after acute stress crosses a tipping point, **(b)** mutagenic stress causing uncontrolled mutagenesis, and **(c)** Inferring phenotypic fitness landscape geometry from sequencing data through selection on standing variation time traces. A power law regime in the mean fitness cost decay reveals the number of fitness-relevant traits. Initial recovery is dominated by selection on the fitness landscape defined by the trait-dependent rate *r*(*z*_1_, *z*_2_). Replicator dynamics, Eq. (1), shifts and contracts the frequency distribution *p*(*z*_1_, *z*_2_, *t*) over time. Here, 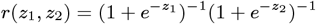 and *p*(*z*_1_, *z*_2_, 0) is Gaussian. **(d)** Calorimetry: the specific heat of a diatomic gas plateaus at values encoding the number of thermally excited degrees of freedom, obeying the law of equipartition [31]. **(e)** Trait counting: the dynamics of the fitness cost for selection on standing variation, Eq. (6), predicts time-dependent plateaus encoding the number of relevant phenotypic dimensions under selection. Data are from Eq. (2) with *r*(*z*_1_, *z*_2_) = 0.1*z*_1_ + 0.0005*z*_2_, *N*_0_(*z*_1_, *z*_2_) = 1 and sampling *z*_*i*_ = 1, 2, …, *N*_*i*_; *N*_1_ = 8000, *N*_2_ = 20000, with *z*_1_, *z*_2_ represented by width and height, respectively.

The answer to this question can be elegantly cast in the language of geometry through the notion of a fitness landscape (see Fig. 1(c), rightmost panel). Concretely, a fitness landscape is the surface obtained by assigning to each trait vector **z** a height equal to its rate *r*(**z**), i.e., the graph of the map **z** ↦ *r*(**z**). Recovery corresponds to an optimization flow on this landscape: because higher-*r* traits recover faster, the trait distribution is progressively reweighted toward higher regions where fitness is greater. As a result, the observed population trajectory reflects both the landscape geometry and the initial trait distribution at *t* = 0.

Identifying experimentally relevant traits **z** that define a fitness landscape *r*(**z**) and the corresponding optimization dynamics calls for data-driven approaches applied to controlled experimental settings. In Fig. 1(c), we present an approach that infers the trait dimensionality of the fitness landscape (rightmost panel) starting from experimental fitness data (leftmost panel). Our key insight is that the two are connected by the recovery trajectory of the fitness *r*(**z**), which we show to exhibit a universal power-law relaxation. Using a mathematical mapping to the equipartition law of thermodynamics (Fig. 1(d)), we show that the recovery provides a quantitative readout of the effective number of traits that determine the fitness landscape (Fig. 1(e)), very much like a calorimetry measurement tells us the number of effective degrees of freedom *n* (e.g. translational, rotational etc.) of a gas whose molar heat capacity is *nR/*2. Once the dimensionality of the fitness landscape is determined, we prove that the resulting mean trait adaptation post-catastrophe is not described by gradient ascent, instead it follows an algorithm known as Levenberg–Marquardt optimization.

Large asexual populations provide a natural testbed for our approach. Their dynamics are normally shaped by slow evolution through the gradual accumulation of mutations [15–17], but their fate is often decided when the environment changes abruptly and survival becomes a race to recover. Such environmental catastrophes can leave behind many competing variants through various routes. One route is ecological: a diverse community can be damaged by acute stress [18–23], which may weaken the ecological interactions [24, 25] or precipitate ecosystem collapse [26, 27] (Fig. 1(a)). A second route is genetic: mutagenic stressors, such as ionizing radiation or genotoxic chemicals, can edit the genetic code and rapidly diversify a previously uniform population [28–30] (Fig. 1(b)).

## SELECTION ON STANDING VARIATION

In the regime of *selection on standing variation* (SSV), strong selection acts on many pre-existing phenotypic variants, while *de novo* variation, e.g., due to mutations, is negligible. Building on this simple insight, we develop a general theory of the evolutionary process that emerges in the wake of catastrophe and compare its predictions with existing experimental measurements in genetic landscapes of antibiotic resistance.

The selection-dominated dynamics of the SSV regime are described by the replicator equation [32],

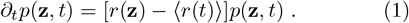

Here, *p*(**z**, *t*) is the average frequency of the trait (phenotype) **z** at time *t* and ⟨*O*(*t*)⟩ ≡ Σ_**z**_ *p*(**z**, *t*)*O*(**z**) denotes the population mean of an observable [33]. Equation (1) can describe a host of recovery processes, where the meaning of *r*(**z**) can range from the Malthusian growth rate under cell division to the stochastic rate for a particular cellstate transition (see Supplementary Information Sec. 9). It describes pure selection dynamics with vanishing mutation, no ecological interactions, and negligible genetic drift. In Supplementary Information Secs. 1–4, we show that the predictions obtained from Eq. (1) are not limited to pure selection: they persist under weak genetic drift [34, 35], weak frequency-dependent selection [32], and weak mutation [36–43]. We also delineate the SSV regime quantitatively, identifying the crossovers at which drift, frequency dependence, or mutation cease to be perturbative, and organize the transition to neighboring evolutionary regimes.

The solution to Eq. (1) is given by

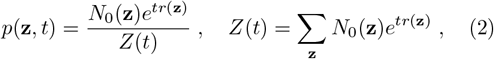

where *N*_0_(**z**) denotes the population’s initial abundances at the onset of catastrophe. In what follows, we elucidate the general behavior of mean observables such as the mean growth rate ⟨*r*(*t*)⟩ and the mean trait ⟨**z**(*t*)⟩.

### SCALE-FREE RECOVERY

As time progresses, if *r*^*∗*^ ≡ *r*(**z**^*∗*^) is the dominant accessible fitness peak, then *p*(**z**, *t*) increasingly concentrates around the corresponding optimum **z**^*∗*^ where the fitness cost *D*(**z**) ≡ *r*^*∗*^ − *r*(**z**) is small. If phenotypic variability exists near **z**^*∗*^, the population finely samples an effectively continuous landscape. In this continuum approximation, Eq. (2) becomes

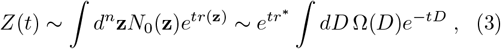

where we introduce the *density of traits*,

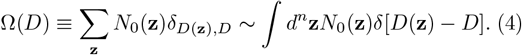

The asymptotic relaxation towards optimality can be characterized systematically by applying Laplace’s method to Eq. (3) in the limit *t* → ∞ (see Methods). Near optimality, where *δz* = |**z** − **z**^*∗*^| is small, the fitness cost is given by *D* ∼ *δz*^*Q*^, and Ω ∼ *D*^*n/Q*−1^ due to *d*^*n*^**z** ∝ *D*^*n/Q*−1^ *dD*. Laplace’s method and the identity 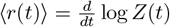 then provide the universal asymptotic form (see Methods),

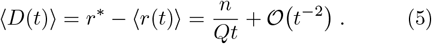

Equation (5) is the first main result: post-catastrophic recovery is slow and scale-free, with a universal, inverse-time decay of the mean fitness cost. This contrasts with the exponential relaxation observed when a beneficial trait fixes in a system with low standing variation. The prefactor depends solely on the local shape parameter *n/Q*, where *n* is the number of non-neutral phenotypic axes near the optimum and *Q* characterizes the local curvature of the peak [44]. For multiple accessible optima, the expansion holds locally: well-separated peaks produce piecewise algebraic relaxation, punctuated by switches between the optima, with an appropriate local value of *n/Q* at each stage (Supplementary Information Sec. 5).

The universality of Eq. (5) admits a simple analogy with equilibrium statistical mechanics. The distribution *p*(**z**, *t*) ∝ *N*_0_(**z**)*e*^−*tD*(**z**)^ is analogous to the equilibrium distribution of a coarse-grained variable **z** at temperature *T* = 1*/t* in the energy landscape *D*(**z**). In this language, Equation (5) is the equipartition law: to leading order, each degree of freedom contributes *T/Q* to the mean energy [45]. Calorimetry relies on equipartition to probe the energy-landscape geometry [31]: intermediate plateaus in specific heat curves probe the effective number of degrees of freedom (Fig. 1(d)). By the same logic, a catastrophe may act as a dynamical probe of the dimensionality of the underlying fitness landscape: *n/Q* can be measured directly through a kinetic fitness assay at multiple time points, agnostic to the underlying phenotype space. In this experiment, the *specific fitness*,

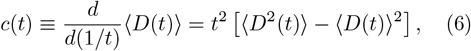

plateaus at *n/Q* in the scale-free regime (Fig. 1(e)). As in gas calorimetry, where plateaus of the specific heat are finite intervals between mode-activation crossovers, the algebraic decay of Eq. (5) and the plateau of *c*(*t*) are observed in an intermediate asymptotic regime *τ*_s_ ≪*t* ≪*τ*_c_ [46]. Here, *τ*_s_, defined by 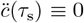, marks the end of the initial transient, while *τ*_c_ ∼ 1*/*Δ*r* marks the breakdown of the continuum approximation, beyond which selection occurs between a small, discrete number of traits separated by a characteristic fitness gap Δ*r* (see Methods for precise form). Independent trait subspaces with additive fitness effects may acquire distinct intermediate asymptotic plateaus, giving rise to the “pyramid” in Fig. 1(e). In Supplementary Information Sec. 6 we show using a scaling theory that this phenomenology extends to weakly-correlated trait subspaces.

In the intermediate asymptotic regime, knowing one of *n* or *Q* determines the other via the measurement of *n/Q*. Generically, *Q* = 1 is expected for directional selection, where **z**^*∗*^ lies at the boundary of the sampled trait space, whereas *Q* = 2 is expected for stabilizing selection, where **z**^*∗*^ lies in its interior [47]. Technical exceptions to this rule are discussed in Methods. Fixing *Q* according to these guidelines and biological insight, the phenotypic dimension *n*, also known as the phenotypic complexity, can be inferred purely from fitness data.

Moreover, the independent growth in the idealized SSV regime implies that Eq. (2) holds at all levels of description: from genotype through mesoscopic phenotypes to complex macroscopic phenotypes. We thus follow the approach in Figure 1(c) to probe the phenotypic dimension in genotypic fitness assays using SSV. We apply this framework to two large experimental datasets in *E. coli* [48, 49] which allows us to identify candidates for the relevant phenotypic axes near optimality.

### PHENOTYPIC DIMENSION INFERENCE

Figure 2 juxtaposes two static assays [48, 49] of antibiotic fitness landscapes in *E. coli*. Static assays measure fitness values across an ensemble of types, yielding Ω(*D*). These data determine the kinetic assay, which tracks a recovering population over time, measuring *Z*(*t*) or equivalently 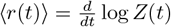. The two assays are linked by the Laplace transform, Eq. (3), analogous to the equivalence of the microcanonical and canonical ensembles in statistical mechanics. For both datasets, the static and kinetic assays are consistent with Eq. (5) and with the common shape parameter *n/Q* = 2. We now show that in each case the local landscape is consistent with *Q* = 1 and *n* = 2.

**FIG. 2.**
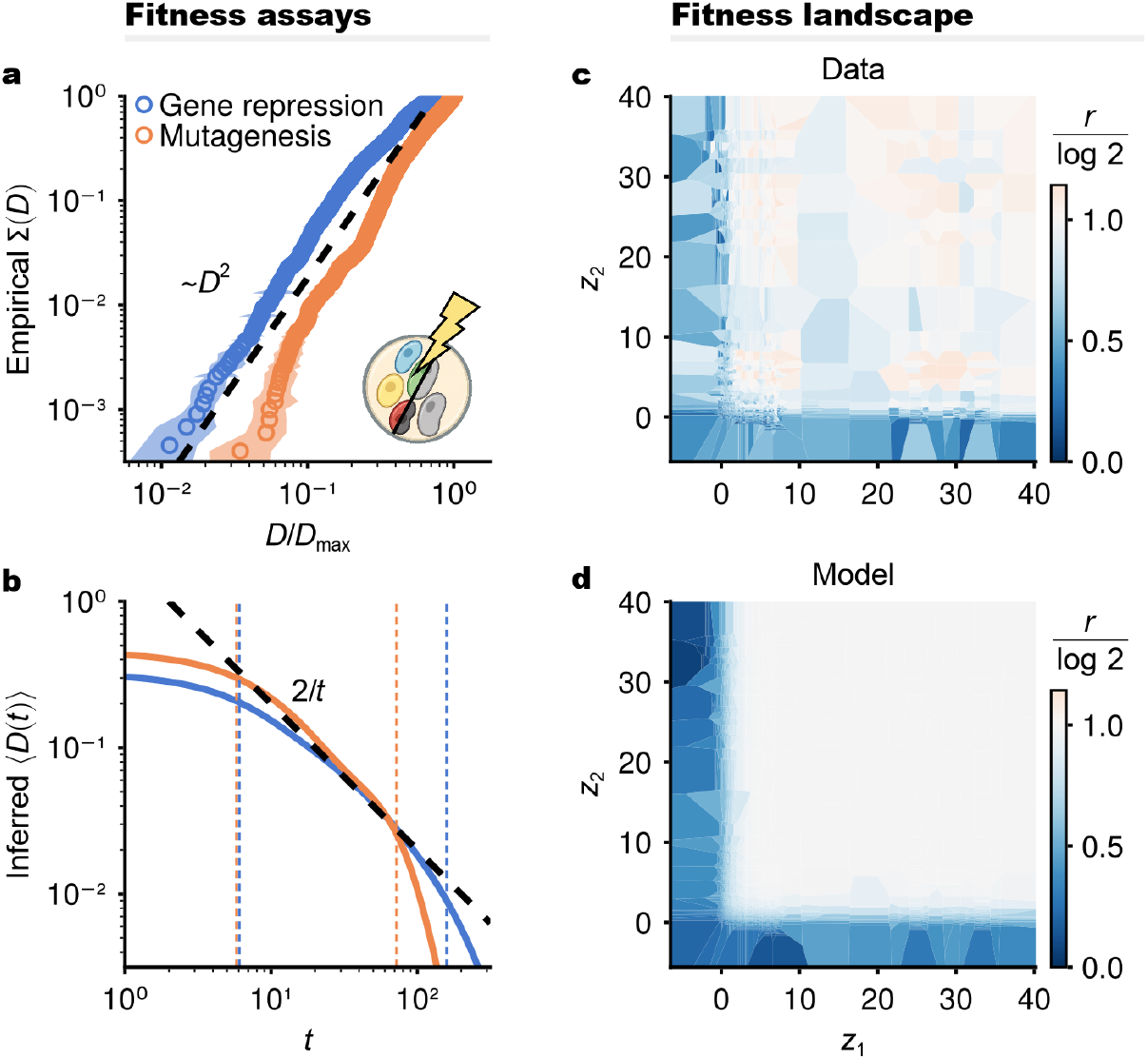
Static fitness assays in *E. coli* measure the phenotypic dimension relevant to evolutionary response. Two complementary assays measure *n/Q*, where *n* is the number of independent traits under selection and *Q* describes the curvature of the fitness peak. Static assays screen a whole library of variants at once, giving a distribution of fitness costs relative to the best variant; the cumulative fraction of variants within a rescaled fitness cost *ρ* = *D/D*_max_ is predicted to scale as Σ(*ρ*) ∼ *ρ*^*n/Q*^. Kinetic assays track a recovering population over time, giving the mean remaining fitness cost ⟨*D*(*t*)⟩ which is predicted to decay as (*n/Q*)*/t* (Eq. (5); the 1*/t* slope is universal). The two are interconvertible by a Laplace transform (Eq. (3)). **(a, b)** Antibiotic fitness landscapes measured statically by Otto et al. [48] via gene-repression (𝒩 = 4464) and Papkou et al. [49] *folA* mutagenesis (𝒩 = 261332); **(a)** the measured distribution (open circles), **(b)** the same data mapped to kinetic form (solid lines). Dashed lines are theory with no fitting parameters; vertical dashed lines mark *τ*_s_ and *τ*_c_, between which the law applies. Both are consistent with *n/Q* = 2, i.e. two-trait landscapes with monotonic peaks. **(c**,**d)** Gene repression assay data and modeling [48] indicate a two-dimensional and monotonic fitness landscape 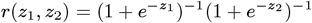.

Figure 2(a-b) reanalyzes static fitness assays under antibiotic stress. The first is a multi-gene double repression assay [48]. Using the CRISPR-i method, the authors imposed variable repression levels *R*_*i*_, *R*_*j*_ ∈ [0, 1] on genes *i, j* from a pool of nine genes, where *R*_*i*_ = 0 denotes unperturbed expression and *R*_*i*_ = 1 complete knockdown. The experiment was found to be well-described by the fitness model,

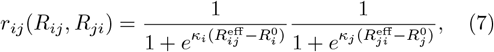

where 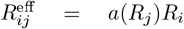 are effective repression levels renormalized by gene-gene interactions *a*(*R*_*j*_). We therefore identify the fitness landscape 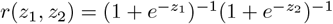 as a function of the effective expression phenotypes 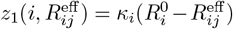 and 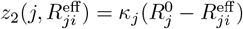 (see Fig. 2(c-d)).

The monotonicity of the landscape implies *Q* = 1, while the two coordinates imply *n* = 2, consistent with Fig. 2(c-d). Furthermore, specific genetic backgrounds correspond to interpretable subspaces of the (*z*_1_, *z*_2_) landscape. By conditioning on repression of gene *i*, we obtain a reduced landscape for which we infer *n/Q* using the standard Clauset–Shalizi–Newman method (CSN), which provides a maximum likelihood estimate over bootstrap-resampled landscapes [50] (see Methods). Then, genetic backgrounds with *n/Q* lower than the wild type correspond to phenotypic subspaces with potentially reduced dimensions. For example, while the full repression dataset yields *n/Q* = 1.9 ± 0.28, conditioning on repression of *gdhA* or *gltB* yields *n/Q* = 1.48 ± 0.27 and 1.76 ± 0.45, respectively. These scalar readouts reflect broad, bimodal bootstrap distributions, arising from the experimental sampling of two distinct regions of the effective landscape *r*(*z*_1_, *z*_2_): an effectively one-dimensional region, where a neutral direction is present, and a two-dimensional region, where it is absent (Supplementary Information Sec. 7). Repeating the analysis of Fig. 2 in these backgrounds reveals that the *gdhA*/*gltB*-conditioned landscapes are predominantly one-dimensional (Extended Figs. 5–6). This is consistent with the known *gdhA*/*gltB* interaction: repression of either gene alone has little fitness effect, whereas only combined repression is lethal [48, 51]. Thus, conditioning on either *gdhA* or *gltB* probes a region of the effective landscape *r*(*z*_1_, *z*_2_) with a neutral direction.

The second assay is a combinatorially-complete, pooled fitness assay of an *E. coli folA* gene segment under trimethoprim antibiotic stress [49]. Using CRISPR-Cas9, the authors generated an *in vivo* library spanning 99.7% of the genotypic variants at nine nucleotide positions encoding amino acid sites 26–28 of dihydrofolate reductase (DHFR), a central metabolic enzyme targeted by trimethoprim. Figure 2(a-b) shows this landscape is consistent with *n/Q* = 2. We argue *Q* = 1, and hence *n* = 2, on kinetic grounds as follows. Enzymatic activity *v*, whose contribution to growth is mediated by metabolic flux [52–55], is captured, in the absence of inhibitor, by the prototypical Michaelis-Menten model

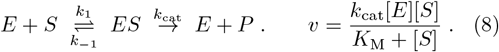

Here, [*E*] and [*S*] are the intracellular enzyme and substrate concentrations and *K*_M_ = (*k* _−1_ + *k*_cat_)*/k*_1_ is the Michaelis constant. Under a competitive inhibitor *I*, such as trimethoprim, *K*_M_ is replaced by 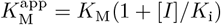, where *K*_i_ is the inhibition constant. The monotonicity of *v* = *v*(*k*_cat_,*K*_M_,*K*_i_,[*E*],[*S*],[*I*]) implies *Q* = 1 near optimality. The readout *n/Q* = 2 in Fig. 2 therefore implies *n* = 2 relevant trait axes. We thus hypothesize the phenotypic landscape *v*(*z*_1_, *z*_2_) = *z*_1_*/*(1 + *z*_2_), where *z*_1_ = *k*_cat_[*E*] and 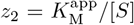 are plausibly distinct axes. This is supported by large-scale enzyme surveys, which find only weak global correlation between *k*_cat_ and *K*_M_ [56], and DHFR mutational studies, which show that nearby activesite substitutions can shift kinetic parameters in various directions [57, 58]. This dimensional reduction recurs in kinetic modeling of DHFR mutations [59, 60], minimal models of drug resistance [61–63], and pharmacodynamic descriptions [61, 64]. More broadly, data-driven models suggest that *n* = 2 collective coordinates can capture functional variation across protein families [65, 66].

Microscopically, the phenotypic dimension *n* = 2 is also supported by the enzyme structure [67] and the topology of the fitness basins [49]. The presence of aspartic acid and glutamic acid at site 27 (D27/E27) is one of the primary distinctions between bacterial and vertebrate DHFRs, respectively, yet the ligand-bound wildtype and D27E structures are nearly identical: although glutamic acid includes an additional methylene group in its side chain, it extends away from the binding site [67]. Consistent with this, Ref. [49] reports that the fitness basin for DHFR variants with either D27 or E27 is smooth and continuous. This is supported by our CSN analysis, which finds the phenotypic dimension conditioned on D27 or E27 to be *n/Q* = 1.92 ± 0.15, 1.82 ± 0.49 respectively, while combining the two conditioned spaces increases it to *n/Q* = 2.57 ± 0.23. The increase reflects reduced constraint in the shared smooth basin and the reduced tendency of mutations at sites 26 and 28 to dramatically affect function.

By contrast, cysteine at site 27 (D27C) leads to a different active site architecture: the conformation of the backbone remains similar, but the cysteinyl sulfur atom accepts a hydrogen bond from T113, bridging the active site and constraining the *α*-helix and *β*-sheet holding C27 and T113, respectively [67]. The thiol group of C27 is critical for substrate protonation, substituting for the function of the carboxyl group present in aspartic and glutamic acid [67]. This binding-site constraint is consistent with selection analysis in Ref. [49] and our reanalysis: D27C variants form a rougher, separate basin with *n/Q* = 1.67 ± 0.31, and site 28 is enriched for leucine after selection relative to D27E or wildtype backgrounds. When conditioned on the presence of L28, the C27 basin increases in average fitness and measured dimension, with *n/Q* = 1.86 ± 0.20. Thus, before site 28 fixes, the dominant phenotypic axis is site 28 itself, whereas after fixation, the smoother basin resembles the D/E27 basin. While this dimensionality readout does not identify molecular phenotypes explicitly, it reveals their conditional structure and provides insight into their microscopic origins.

The above analyses go beyond previous attempts to measure the phenotypic dimension using effective phenotypic evolution models [44, 68, 69]. The SSV regime makes phenotypic dimension measurable from the idealized recovery dynamics alone, agnostic to the unknown genotype–phenotype and phenotype–fitness maps required in mutation-limited settings [17, 44, 70–72]. Moreover, it enables a direct comparison between information content and phenotypic dimension. The information content of a library of 𝒩 variants is *L* ≡ log_2_ 𝒩 [73].

The landscapes of Refs. [48, 49] analyzed in Fig. 2(a-b) have information content *L* ≃ 12, 18, respectively. A non-redundant sampling of a phenotypic space of dimension *n*, characteristic size *X*, and mesh size *a* gives 𝒩 ∼ (*X/a*)^*n*^, or equivalently *L* ∼ log(*X/a*)*n*. Our analysis yields *L/n* = 6, 9 for the respective landscapes, consistent with limited (subexponential) redundancy of traits in the sampled space. This is also consistent with the phenotypic map (*i, j, R*_*i*_, *R*_*j*_) ⟼ *z*_*k*_ found for Ref. [48], which displays strong phenotypic epistasis, whereby the effect of a mutation depends strongly on its background, and pleiotropy, whereby a mutation can affect several traits simultaneously [44, 74]. Despite the lack of direct access to the genotype–phenotype map, our measurement of the phenotypic dimension for Ref. [49] likewise supports the presence of these effects.

### NON-GRADIENT OPTIMIZATION

The inverse-time law in Eq. (5) constrains the scalar recovery. A complementary question is therefore how the dominant trait itself evolves in phenotype space. Let **z** = **z**(*t*) denote the highest-frequency trait, around which *p*(**z**, *t*) concentrates. In the SSV regime this trait climbs toward higher fitness, but not generally by steepest ascent (Fig. 3(a)). Maximizing the distribution in Eq. (2) implies

**FIG. 3.**
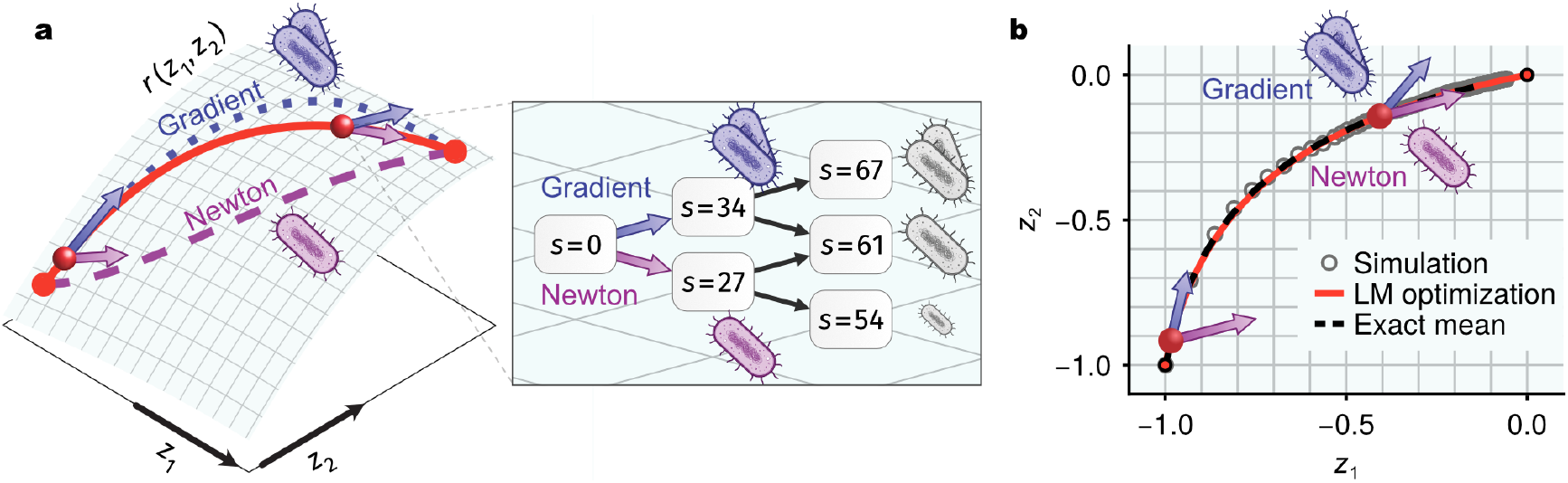
Selection on standing variation follows curvature-sensitive trajectories on fitness landscapes. **(a)** Gradient ascent, Newton’s method, and Levenberg-Marquardt dynamics on the landscape 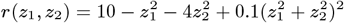, and the Levenberg-Marquardt dynamics which interpolates between them. The Levenberg-Marquardt dynamics adaptively selects between the steepest ascent (gradient, blue) and a curvature-sensitive ascent (Newton, purple), with directions shown by arrows. While gradient flow is preferable on the steep fitness slopes, Newton’s method is preferable near the peak where some directions may be shallow. Newton’s method reweights the diminishing fitness gradient and takes shorter paths in anisotropic landscapes. Inset: Discretization of the trajectory about **z**_0_ = (− 0.42, − 0.14) as a selection “tournament” (lattice spacing *a* = 5 × 10^−3^). The winner of each round is determined by the optimization rule and the selection coefficient *s*(**z**) = [*r*(**z**) − *r*(**z**_0_)]*/κ*_max_*a*^2^ where *κ*_max_ is the largest eigenvalue of − *H*_*ij*_ (**z**^*∗*^) (rounded to the nearest integer for visualization). The gradient rule (blue) chooses the high-*s* successor, whereas the Newton rule (purple) considers the next generation and may locally prefer the less fit successor. **(b)** Highest-frequency trait trajectories **z**(*t*) on the fitness landscape in (a). Stochastic birth-death simulations (circles) are compared with the Levenberg-Marquardt dynamics (Eq. (10)) and the exact solution (Eq. (2); lines). For sufficiently large and peaked initial populations, ⟨**z**(*t*)⟩ converges to the Levenberg-Marquardt trajectory.

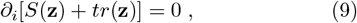

where *S*(**z**) ≡ log *N*_0_(**z**). Differentiating Eq. (9) with respect to time yields

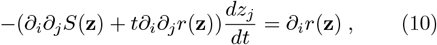

where summation over repeated indices is implied. The solution **z**(*t*) to Eq. (10) recovers the long-time asymptotics of Eq. (5). Moreover, ⟨**z**(*t*)⟩ ∼ **z**(*t*) when *N*_0_(**z**) is sufficiently peaked around an initial trait **z**_0_, as verified by the stochastic simulations in Fig. 3(b) and analytically in the Supplementary Information Sec. 8.

Equation (10) resembles a gradient ascent modified with a time-dependent friction tensor *M*_*ij*_(**z**, *t*) = −*∂*_*i*_*∂*_*j*_[*S*(**z**) + *tr*(**z**)]. It can be recast as a function of *s* = log *t* as

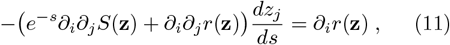

which can be identified as the continuous-time limit of the Levenberg-Marquardt algorithm, a regularized version of Newton’s method [75–79]. As demonstrated in Fig. 3(a) for *n* = 2, the Levenberg-Marquardt trajectory initially follows steepest ascent. As time proceeds, however, the motion becomes increasingly sensitive to landscape curvature through the fitness Hessian *H*_*ij*_ ≡ *∂*_*i*_*∂*_*j*_*r*(**z**), and approaches Newton’s method near the optimum. For *n* = 1, this affects only trajectory speed; for *n* > 1, it changes the path itself.

To illustrate this, we discretize Eq. (10) on a square lattice of spacing *a*. The fitness gradient on the right-hand side of Eq. (10) becomes *∂*_*i*_*r*(**z**) → *r*(**z** + **e**_*i*_) − *r*(**z**), while the fitness Hessian becomes *H*_*ij*_(**z**) → *r*(**z**+**e**_*i*_ +**e**_*j*_) − *r*(**z**+**e**_*i*_) − *r*(**z**+**e**_*j*_)+*r*(**z**), where (**e**_*i*_)_*j*_ = *aδ*_*ij*_. Figure 3(a) then recasts the discrete-time update as a selection tournament. Under the gradient rule, the preferred successor *z*_*i*_ is the one with the larger pairwise fitness gain *∂*_*i*_*r*. Under the Newton rule, the preferred successor depends on three-point comparisons through − (*H*^−1^)_*ij*_*∂*_*j*_*r*, and can therefore locally favor the less fit successor. Strikingly, this remains the case even if the fitter trait has fitter successors.

Predicting post-catastrophic trajectories thus requires more than pairwise fitness differences: three-point comparisons are needed to evaluate local curvature. The full Levenberg-Marquardt trajectory interpolates smoothly between these two schemes and takes progressively greater fitness risks. Near the fitness peak, selection therefore becomes biased against greediness, preferentially exploring locally shallower phenotypic directions—a property known in optimization theory as “optimism”. Thus, from an optimization perspective, post-catastrophic selection on standing variation is optimistic [80].

## ACKNOWLEDGEMENTS

This research was partly supported by the National Science Foundation through the Physics Frontier Center for Living Systems (PHY2317138) as well as NSF (DMS-2235451) and Simons Foundation (MPS-NITMB-00005320) to the NSF-Simons National Institute for Theory and Mathematics in Biology (NITMB). V. V. is a Chan Zuckerberg Biohub Chicago Investigator. O.G. acknowledges support from the Leinweber Institute for Theoretical Physics, the Center for Living Systems at The University of Chicago and a MRSEC-funded Kadanoff–Rice fellowship from The University of Chicago Materials Research Science and Engineering Center, which is funded by NSF (DMR-2011854).

## METHODS

### Laplace’s method

Laplace’s method is a systematic asymptotic expansion of Laplace integrals about the dominant rate. For the partition function *Z*(*t*), inserting Ω(*D*) = Ω_0_*D*^*n/Q*−1^ + 𝒪 (*D*^*n/Q*^) into Eq. (3) yields

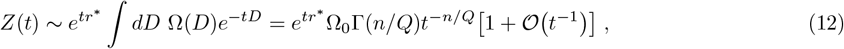

which provides Eq. (5) after taking the logarithmic derivative. The geometric interpretation of Laplace’s method as a locally smooth approximation of the fitness landscape is more apparent by directly expanding the multidimensional integral in Eq. (3). With *δz* = |*δ***z**| and *δ***z** ≡ **z** − **z**^*∗*^, the leading-order fitness cost can be expanded as

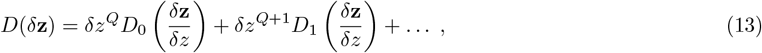

where *Q* > 0. Substituting Eq. (13) into Eq. (3) and applying the change of variables *δ***z** = *t*^−1*/Q*^***ξ***, we obtain for the regular, smooth cases described below,

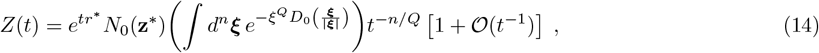

where the integration over ***ξ*** is constrained to the sampled sector of the phenotype space relative to **z**^*∗*^. Comparing with Eq. (12) determines 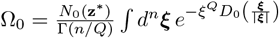. The 𝒪 (*t*^−1^) correction in Eq. (14) is obtained for smooth landscapes with a generic interior maximum (*Q* = 2) or boundary vertex maximum (*Q* = 1). It is likewise obtained for a maximum in the interior of an *m*-dimensional boundary. In that case, one may choose *n* − *m* coordinates normal to the boundary and *m* coordinates tangent to it. Then, the density of traits has *m* quadratic contributions and *n* − *m* linear contributions, again reproducing the 𝒪 (*t*^−1^) correction. In the most general case of a nonsmooth landscape, the correction can remain 𝒪 (*t*^−1*/Q*^), leading to a 𝒪 (*t*^−1−1*/Q*^) correction in Eq. (5). Smooth but degenerate maxima with *Q* > 2 provide another technical exception, with fractional subleading powers even when the landscape is differentiable.

An analogous calculation shows that the population average ⟨*O*(*t*)⟩ of an arbitrary smooth observable *O*(**z**) in an arbitrary landscape is given by

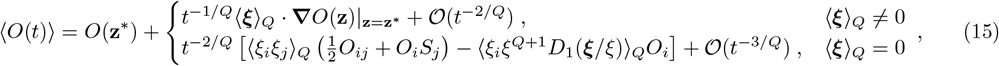

where *O*_*i*_ ≡ *∂*_*i*_*O*|_**z**=**z**_*\**, *O*_*ij*_ ≡ *∂*_*i*_*∂*_*j*_*O*|_**z**=**z**_*\*, S*_*i*_ ≡ *∂*_*i*_ log *N*_0_(**z**)|_**z**=**z**_*\**, *ξ* = |***ξ***|, and

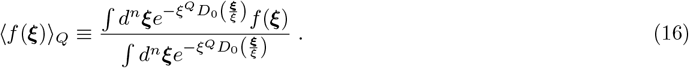

The expansions in Eq. (12) and Eq. (15) indicate that the power law regime generically depends on *N*_0_(**z**) and the observable *O*(**z**). Indeed, in the most general case, the integral in Eq. (15) is sensitive to the local geometry about **z**^*∗*^. Like the correction in Eq. (14), the leading decay in Eq. (15) generically becomes 𝒪 (*t*^−1^) in the regular cases, both in boundary (*Q* = 1, ⟨***ξ***⟩_1_ ≠ 0) and interior (*Q* = 2, ⟨***ξ***⟩_2_ = 0) cases. Fractional powers arise in nonregular exceptions, including nonsmooth landscapes and smooth but degenerate maxima.

### Trait counting and intermediate asymptotics

The early *τ*_s_ and late *τ*_c_ timescales defining the intermediate asymptotic regime *τ*_s_ ≪ *t* ≪ *τ*_c_ correspond to the initial and final inflection points of the specific fitness *c*(*t*), between which a *c* ∼ *n/Q* plateau is obtained.

Fluctuations in real data may cause numerous inflection points in *c*(*t*), despite a single equipartition plateau. The first inflection point *τ*_s_, can be found by solving 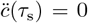 numerically. For a characteristic fitness gap Δ*r* near the optimum *r* = *r*^*∗*^, the final inflection point *τ*_c_ ∼ 1*/*Δ*r* can be determined via the late-time expansion 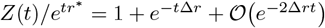.

A more accurate determination of *τ*_c_ employs a criterion from the theory of Bose-Einstein condensates [81, 82]. Splitting the partition function into discrete and continuous parts

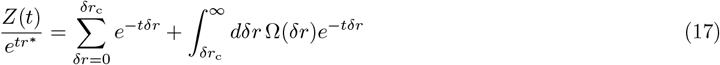

The time *τ*_c_ is then the crossover point where the discrete and continuous distributions balance,

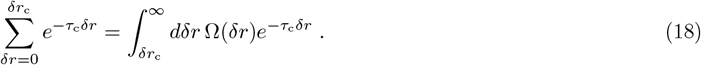

By Laplace’s method, the right-hand side satisfies 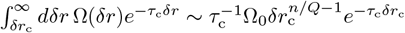 and vanishes if *τ*_c_ → ∞. In contrast, the left-hand side approaches unity. Therefore, *δr*_c_ must be small enough that 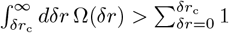 which guarantees that Eq. (18) has a solution. In Fig. 2(b) we solve Eq. (18) numerically with *δr*_c_ = Δ*r*.

The above construction explains how several plateaus can appear in a single readout of *c*(*t*). Consider trait subspaces **z** = (**z**_1_ **z**_*m*_) whose fitness effects are additive over the relevant range, *r* (**z**) = Σ _*α*_ *r*_*α*_ (**z**_*α*_), and whose initial distributions are factorizable, *p*_0_(**z**) = Π_*α*_ *p*_0,*α*_(**z**_*α*_). Then, Eq. (6) for subspace *α*, implies

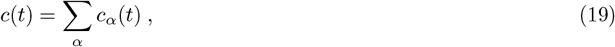

where we denote the contribution by subspace *α* using a subscript. Each subspace thus has its own intermediate asymptotic regime *τ*_s,*α*_ ≪ *t* ≪ *τ*_c,*α*_ in which *c*_*α*_(*t*) ≃ *n*_*α*_*/Q*_*α*_. The specific fitness measures the sum of the contributions under SSV at that time (Fig. 1(e)), much akin to calorimetry, where different mechanical modes are thermally activated over different temperature regimes (Fig. 1(d)). In Supplementary Information Sec. 6, we show that the above analysis extends to weakly correlated subspaces with non-additive effects using a scaling theory.

### Dimension estimation by Clauset–Shalizi–Newman power-law fitting

Given experimental measurements 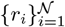 of a fitness landscape over types *i*, we define the fitness cost *D*_*i*_ ≡ *r*^*∗*^ − *r*_*i*_, where *r*^*∗*^ ≡ max_*j*_ *r*_*j*_. In the scale-free regime, the integrated density of traits satisfies

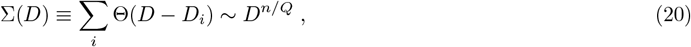

thus providing a readout of the shape parameter *n/Q* by the predicted linear relationship 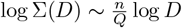.

To provide a principled error estimate for this readout, we use the CSN power-law maximum-likelihood procedure [50, 83]. If the power law Σ(*D*) ∼ *D*^*n/Q*^ holds up to a cutoff scale *D*_c_, the maximum likelihood estimator is

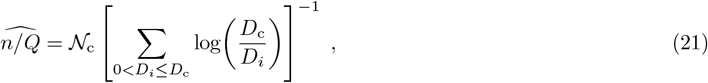

where *𝒩*_c_ ≡ Σ_0*<Di≤D*c_ and the cutoff scale *D*_c_ is selected to minimize the Kolmogorov–Smirnov statistic,

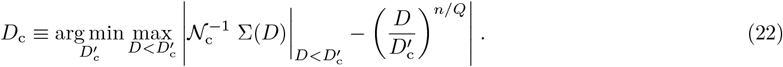

The uncertainty in *D*_c_ is quantified by bootstrapping with many resamples of the data. Here, we use bootstrapping with 10^4^ resamples and report the bootstrap average and standard deviation. This gives an objective, dataset-agnostic statistical procedure to estimate *n/Q*. Because reduced landscapes conditioned on specific genetic backgrounds contain fewer points, their estimates are more sensitive to the inferred cutoff *D*_c_, which sets the dynamic range of the data that contributes to the estimate. In these cases, the bootstrap distribution itself is informative; broad or multimodal distributions are inspected separately in Supplementary Information Sec. 7.

### Stochastic simulations

To show that our results deriving from Eq. (2) are robust to demographic noise, we conduct a stochastic simulation of a population growing with trait-dependent birth rates *r*(**z**) given by the fitness landscape of Fig. 3(a) and vanishing death rates, resulting in the trajectory shown in Fig. 3(b). We use a standard Gillespie *τ* -leaping scheme [84] with *τ* = 0.1 and couple the population to a chemostat by periodically diluting it back to its initial population *Z*(0) = 10^6^ every Δ*t* = 1 time units, congruent with standard experimental setups in microbial evolution [85]. The initial population *N*_0_(**z**) is an isotropic Gaussian of variance *σ*^2^ = 1*/*24, ensuring convergence to Eq. (10) (Supplementary Information Sec. 9). An empirical population drawn from the distribution *N*_0_(**z**) has negligible probability of providing fine sampling near the optimum. We therefore utilize an importance-biased Monte Carlo technique [86, 87] by simulating from an initially uniform distribution over the traits and estimating the average trait averaged over trajectories via

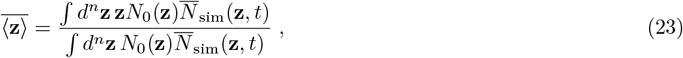

where *N*_sim_(**z**, *t*) is the simulated population. This estimator is asymptotically exact for large populations, where 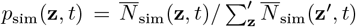 converges to the replicator solution, Eq. (2). Further details are given in Supplementary Information Sec. 9.

